# VarSight: Prioritizing Clinically Reported Variants with Binary Classification Algorithms

**DOI:** 10.1101/532440

**Authors:** James M. Holt, Brandon Wilk, Camille L. Birch, Donna M. Brown, Manavalan Gajapathy, Alexander C. Moss, Nadiya Sosonkina, Melissa A. Wilk, Julie A. Anderson, Jeremy M. Harris, Jacob M. Kelly, Fariba Shaterferdosian, Angelina E. Uno-Antonison, Arthur Weborg, Undiagnosed Diseases Network, Elizabeth A. Worthey

## Abstract

**Motivation:** In genomic medicine for rare disease patients, the primary goal is to identify one or more variants that cause their disease. Typically, this is done through filtering and then prioritization of variants for manual curation. However, prioritization of variants in rare disease patients remains a challenging task due to the high degree of variability in phenotype presentation and molecular source of disease. Thus, methods that can identify and/or prioritize variants to be clinically reported in the presence of such variability are of critical importance.

**Results:** We tested the application of classification algorithms that ingest variant annotations along with phenotype information for predicting whether a variant will ultimately be clinically reported and returned to a patient. To test the classifiers, we performed a retrospective study on variants that were clinically reported to 237 patients in the Undiagnosed Diseases Network.
We treated the classifiers as variant prioritization systems and compared them to four variant prioritization algorithms and two single-measure controls. We showed that these classifiers outperformed the other methods with the best classifiers ranking 72% of all reported variants and 94% of reported pathogenic variants in the top 20.

**Availability:** The scripts used to generate results presented in this paper are available at https://github.com/HudsonAlpha/VarSight release v1.1.

## 1 Introduction

Genome and exome sequencing are both currently being used as molecular diagnostic tools for patients with rare, undiagnosed diseases (Ramoni *et al.*, 2017; Bagnall *et al.*, 2018; Sweeney *et al.*, 2018). Typically, these technologies are applied clinically following workflows consisting of blood draw, sequencing, alignment, variant calling, variant annotation, variant filtering, and variant prioritization (Worthey, 2017; Roy *et al.*, 2018). Then, clinical analysts usually perform the more manual processes of inspecting and then clinically reporting variants based on the known set of patient phenotypes.

In general, commonly used pipelines exist for the steps from sequencing through variant calling (Rehm *et al.*, 2013; Cornish *et al.*, 2015). Despite differences in performance, most of these standards ingest the same information to create a list of variants from sequencing data. In contrast, methods for variant annotation and/or variant filtering are quite diverse (Wang *et al.*, 2010; Hu *et al.*, 2013; Jger *et al.*, 2014; Desvignes *et al.*, 2018). These methods use a wide range of input sources including but not limited to population allele frequencies (Lek *et al.*, 2016), conservation scores (Cooper *et al.*, 2005; Siepel *et al.*, 2006; Petrovski *et al.*, 2013), haploinsufficiency scores (Huang *et al.*, 2010; Steinberg *et al.*, 2015), deleteriousness scores (Steinberg *et al.*, 2015; Rentzsch *et al.*, 2018), transcript impact scores (Kumar *et al.*, 2009; Choi, 2012; Adzhubei *et al.*, 2013; Dong *et al.*, 2014; Jian *et al.*, 2014), and previously associated disease annotation (Stenson *et al.*, 2003; Hamosh *et al.*, 2005; Landrum *et al.*, 2015). Variant prioritization is also quite diverse with some methods relying on the variant annotations to prioritize variants (Hu *et al.*, 2013) and some relying on patient phenotype to rank the variants (Khler *et al.*, 2009; Yang *et al.*, 2015; Rao *et al.*, 2018; Wilk *et al.*, 2018). There are also methods which combine both variant annotations and phenotype score to rank the variants (Singleton *et al.*, 2014; Zemojtel *et al.*, 2014; Smedley *et al.*, 2015a; Boudellioua *et al.*, 2019), a selection of which are benchmarked on the same simulated datasets in Smedley et al. (2015b).

Given a prioritized list of variants, analysts manually inspect those variants and curate a list of variants to ultimately report to the ordering physician. Unfortunately, manual curation is a time consuming process where analysts must inspect each variant while maintaining a mental picture of the patient’s phenotype. Bick *et al.*, 2017 report an average of 600 variants per case analyzed by two people (one analyst and one director) over three hours, meaning a thoughput of ≈100 variants per man-hour. If causative variants can be identified earlier due to a high rank from prioritization, it’s possible that the full filtered variant list can be short-circuited, reducing the total number of variants reviewed and therefor the time to analyze a case. Additionally, accurate prioritization is a step towards the ultimate goal of automatic identification of rare variants that may cause a patient’s primary phenotypes.

One of the issues with previously published ranking methods is that they were primarily tested on simulated datasets with known, single-gene, pathogenic variants injected into real or simulated background genomic datasets. Additionally, when phenotype terms were used, they tended to use all available phenotype terms paired with the simulated disease with a few (typically 2-3) noisy terms added or removed. In practice, rare disease patient data is often far noisier due to a wide variety of reasons.

In this paper, we focus on real patient data from the multi-site collaboration of the Undiagnosed Diseases Network (UDN) (Ramoni *et al.*, 2017). Patients accepted into the UDN are believed to have rare, undiagnosed diseases of genetic origin. Because the UDN is not focused on a single particular disease, the patient population has a diverse range of phenotypes represented. Additionally, the phenotypes associated to an individual patient can be quite noisy for a variety of reasons: multiple genetic diseases, phenotype collection differences, and/or unrelated non-genetic diseases (such as age- or pathogen-related disease). Because the UDN is a research collaboration, there is also variability in reported variants that range in pathogenicity from “variant of uncertain significance” (VUS) through “pathogenic” as defined by the ACMG guidelines (Richards *et al.*, 2015). The summation of this real-world variation means that accurately prioritizing variants is challenging due to noise and variation in phenotype inputs and variation in pathogenicity of reported variant outputs.

## 2 Approach

We tested the application of classification algorithms for identifying clinically reported variants in real world patient in two ways: 1) predicting whether a variant observed by an analyst would be clinically reported and 2) prioritizing all variants seen by the clinical analysts. In particular, we focused our analyses on real patients with a diverse collection of rare, undiagnosed diseases that were admitted to the Undiagnosed Diseases Network (UDN) (Ramoni *et al.*, 2017). We limited our patients to those who received whole genome sequencing and received at least one primary variant (i.e. not secondary or incidental) on their clinical report. We extracted data directly from the same annotation and filtering tool used by the analysts in order to replicate their data view of each variant in a patient. Additionally, we incorporated phenotype information into the models using two scoring systems that are based on ranking genes by their association to a set of patient phenotypes. Finally, each variant was either labeled as “returned” or “not returned” depending on whether it was ultimately reported back to the clinical site.

Given the above variant information, we split the data into training and testing sets for measuring the performance of classifiers to predict whether a variant would be clinically reported or not. We tested four classifiers that are readily available in the *sklearn* (Pedregosa *et al.*, 2011) and *imblearn* (Lematre *et al.*, 2017) Python modules. Of note, our focus was not on picking the “best” classifier, but rather on analyzing their overall ability to handle the noise of real-world patient cases from the UDN.

Each classifier calculated probabilities of a variant belonging to the “returned” class, allowing us to measure their performance as both a classifier and a prioritization/ranking system. After tuning each classifier, we generated summaries of the performance of each method from both a binary classification perspective and a variant prioritization perspective. Additionally, we tested four publicly available variant prioritization algorithms and two single-value ranking methods for comparison. All of the scripts to train classifiers, test classifiers, and format results are contained in the VarSight repository.

## 3 Methods

### 3.1 Data sources

All samples were selected from the cohort of Undiagnosed Diseases Network (UDN) (Ramoni *et al.*, 2017) genome sequencing samples that were sequenced at HudsonAlpha Institute for Biotechnology. In short, the UDN accepts patients with rare, undiagnosed diseases that are believed to have a genetic origin. The UDN is not restricted to a particular disease, so there are a diverse set of diseases and phenotypes represented across the whole population. The phenotypes annotated to a patient are also noisy compared to simulated datasets for a variety of reasons including: 1) patients may have multiple genetic diseases, 2) phenotype collection is done at seven different clinical sites leading to slightly different standards of collection, 3) patients may exhibit more or fewer phenotypes than are associated with the classic disease presentation, and 4) patients may have phenotypes of non-genetic origin such as age- or pathogen-related phenotypes. For more details on the UDN, refer to Ramoni *et al.*, 2017.

DNA for these UDN patients was prepared from blood samples (with few exceptions) and sequenced via standard operation protocols for use as a Laboratory-Developed Test (LDT) in the HAIB CAP/CLIA lab. The analyses presented in this paper are based on data that is or will be deposited in the dbGaP database under dbGaP accession phs001232.v1.p1 by the UDN.

### 3.2 Alignment and variant calling

After sequencing, we followed GATK best practices (DePristo *et al.*, 2011) to align to the GRCh37 human reference genome with BWA-mem (Li, 2013). Aligned sequences were processed via GATK for base quality score recalibration, indel realignment, and duplicate removal. Finally, SNV and indel variants were joint genotyped, again according to GATK best practices (DePristo *et al.*, 2011). The end result of this pipeline is one Variant Call Format (VCF) file per patient sample. This collection of VCF files is used in the following sections.

### 3.3 Variant annotation and filtering

After VCF generation, the clinical analysts followed various published recommendations (e.g. Worthey, 2017; Roy *et al.*, 2018) to annotate and filter variants from proband samples. For variant annotation and filtering, we used the same tool that our analysts used during their initial analyses. The tool, Codicem (Envision, 2018), loads patient variants from a VCF and annotates the variants with over fifty annotations that the analysts can use to interpret pathogenicity. These annotations include: variant level annotations such as CADD (Rentzsch *et al.*, 2018), conservation scores (Cooper *et al.*, 2005; Siepel *et al.*, 2006), and population frequencies (Lek *et al.*, 2016); gene level annotations such as haploinsufficiency scores (Huang *et al.*, 2010; Steinberg *et al.*, 2015), intolerance scores (Petrovski *et al.*, 2013), and disease associations (Stenson *et al.*, 2003; Hamosh *et al.*, 2005; Landrum *et al.*, 2015); and transcript level annotations such as protein change scores (Kumar *et al.*, 2009; Choi, 2012; Adzhubei *et al.*, 2013; Dong *et al.*, 2014) and splice site impact scores (Jian *et al.*, 2014). Additionally, if the variant has been previously curated in another patient through HGMD or ClinVar (Stenson *et al.*, 2003; Landrum *et al.*, 2015), those annotations are also made available to the analysts.

Codicem also performs filtering for the analysts to reduce the number of variants that are viewed through a standard clinical analysis. We used the latest version of the primary clinical filter for rare disease variants to replicate the standard filtering process for patients in the UDN. In short, the filter requires the following for a variant to pass through the clinical filter: sufficient total read depth, sufficient alternate read depth, low population frequency, at least one predicted effect on a transcript, at least one gene-disease association, and to not be a known, common false-positive from sequencing. In general, the filter reduces the number of variants from the order of millions to hundreds (anecdotally, roughly 200-400 variants per proband after filtering). For the specific details on the filter used, please refer to Supplementary Documents.

### 3.4 Phenotype annotation

The Codicem annotations are all agnostic of the patient phenotype. As noted earlier, we expect these patient phenotypes to be noisy when compared to simulated datasets due to the variety and complexity of diseases, phenotypes, and genetic heritage tied to UDN patients. Additionally, we made no effort to alter or condense the set of phenotypes provided by the corresponding clinical sites. In order to incorporate patient phenotype information, we used two distinct methods to rank genes based on the Human Phenotype Ontology (HPO) (Khler *et al.*, 2018). We then annotated each variant with the best scores from their corresponding gene(s).

The first method uses phenotype-to-gene annotations provided by the HPO to calculate a cosine score (Khler, 2017) between the patient’s phenotypes and each gene. Given P terms in the HPO, this method builds a binary, *P*-dimensional vector for each patient such that only the phenotype terms and all ancestral phenotype terms tied to the patient are set to 1. Similarly, a *P*-dimensional vector for each gene is built using the phenotype-to-gene annotations. Then, the cosine of the angle between the patient vector and each gene vector is calculated as a representation of similarity. This method tends to be more conservative because it relies solely on curated annotations from the HPO.

The second method, an internally-developed tool called PyxisMap (Wilk *et al.*, 2018), uses the same phenotype-to-gene annotations from the HPO, but adds in automatically text-mined annotations from NCBI’s PubTator (Wei *et al.*, 2013) and performs a Random-Walk with Restart (Page *et al.*, 1999) on the ontology graph structure. The PyxisMap method has the added benefit of incorporating gene-phenotype connections from recent papers that have not been manually curated into the HPO, but it also tends to make more spurious connections due to the imprecision of the text-mining from PubTator. Each method generates a single numerical feature that is used in the following analyses.

### 3.5 Patient selection

In the clinical analysis, each patient was fully analyzed by one director and one analyst. After the initial analysis, the full team of directors and analysts review flagged variants and determine their reported pathogenicity. In our analysis, we focused on variants that were clinically reported as “primary”, meaning the team of analysts believed the variant to be directly related to the patient’s phenotype. Note that secondary and/or incidental findings are specifically not included in this list. The team of analysts assigned each primary variant a classification from variant of uncertain significance (VUS), likely pathogenic, or pathogenic adhering to the recommendations in the ACMG guidelines for variant classification (Richards *et al.*, 2015).

We required the following for each proband sample included in our analyses: 1) at least one clinically reported primary variant that came through the primary clinical filter (i.e. it was not found through some other targeted search) and 2) a set of phenotypes annotated with Human Phenotype Ontology (Khler *et al.*, 2018) terms using the Phenotips software (Girdea *et al.*, 2013). At the time of writing, this amounted to 378 primary-reported variants and 87819 unreported variants spanning a total of 237 proband samples.

### 3.6 Feature Selection

For the purposes of classification, all annotations needed to be cleaned and stored as numerical features. For single-value numerical annotations (e.g. float values like CADD or GERP), we simply copied the annotation over as a single value feature. Missing annotations were assigned a default value that was outside the expected value range for that feature. Additionally, these default values were always on the less impactful side of the spectrum (e.g. a default conservation score would err on the side of not being conserved). The one exception to this rule was for variant allele frequencies where a variant absent from the database was considered to have an allele frequency of 0.0. For multi-value numerical annotations, we reduced the values (using minimum or maximum) to a single value corresponding to the “worst” value (i.e. most deleterious value, most conserved value, etc.) that was used as the feature.

For categorical data, we relied on bin-count encoding to store the features. We chose to bin-count because there are many annotations where multiple categorical labels may be present at different quantities. For example, a single ClinVar variant may have multiple entries where different sites have selected different levels of pathogenicity. In this situation, we desired to capture not only the categorical label as a feature, but also the number of times that label occurred in the annotations.

After converting all annotations to numerical features or multiple bin-count encoded features, we had a total of 95 features per variant. We then pruned down to only the top 20 features using univariate feature selection (specifically the `SelectKBest` method of *sklearn* (Pedregosa *et al.*, 2011)). This method evaluates how well an individual feature performs as a classifier and keeps only the top 20 features for the full classifiers. Table 1 shows the list of retained features ordered by feature importance after training. Feature importance was derived from the random forest classifiers which automatically report how important each feature was for classification. The entire set of annotations along with descriptions of how each was processed prior to feature selection are detailed in the Supplementary Documents.

**Table 1:** Feature selection. This table shows the top 20 features that were used to train the classifiers ordered from most important to least important. After training, the two random forest classifiers report the importance of each feature in the classifier (total is 1.00 per classifier). We average the two importance values, and order them from most to least important. Feature labels with an represent a single category of a multi-category feature (i.e. “HGMD assessment type_DM” means the “DM” bin-count feature from the “HGMD assessment type” annotation in Codicem).

| Feature label | RF(sklearn) | BRF(imblearn) |
| --- | --- | --- |
| HPO-cosine | 0.2895 | 0.2471 |
| PyxisMap | 0.2207 | 0.2079 |
| CADD Scaled | 0.1031 | 0.1007 |
| phylop100 conservation | 0.0712 | 0.0817 |
| phylop conservation | 0.0641 | 0.0810 |
| phastcon100 conservation | 0.0572 | 0.0628 |
| GERP rsScore | 0.0357 | 0.0416 |
| HGMD assessment type_DM | 0.0373 | 0.0344 |
| HGMD association confidence_High | 0.0309 | 0.0311 |
| Gnomad Genome total allele count | 0.0192 | 0.0322 |
| ClinVar Classification_Pathogenic | 0.0228 | 0.0200 |
| ADA Boost Splice Prediction | 0.0081 | 0.0109 |
| Random Forest Splice Prediction | 0.0077 | 0.0105 |
| Meta Svm Prediction_D | 0.0088 | 0.0092 |
| PolyPhen HV Prediction_D | 0.0075 | 0.0071 |
| Effects_Premature stop | 0.0049 | 0.0057 |
| SIFT Prediction_D | 0.0026 | 0.0056 |
| PolyPhen HD Prediction_D | 0.0025 | 0.0049 |
| Effects_Possible splicing modifier | 0.0029 | 0.0035 |
| ClinVar Classification_Likely Pathogenic | 0.0034 | 0.0020 |

### 3.7 Classifier training and tuning

As noted earlier, there are generally hundreds of variants per proband that pass the filter, but only a few are ever clinically reported. Across all 237 proband samples, there were a total of 378 clinically reported variants and another 87819 variants that were seen but not reported. As a result, there is a major imbalance in the number of true positives (variants clinically reported) and true negatives (variants seen, but not clinically reported).

We split the data into training and test sets on a per-proband basis with the primary goal of roughly balancing the total number of true positives in each set. Additionally, the cases were assigned to a particular set by chronological order of analysis in order to reduce any chronological biases that may be introduced by expanding scientific knowledge (i.e. there are roughly equal proportions of “early” or “late” proband samples from the UDN in each set). In the training set, there were a total of 189 returned variants and 44593 not returned variants spanning 120 different probands. In the test set, there were a total of 189 returned variants and 43226 not returned variants spanning 117 different probands. In our results, the returned test variants are further stratified by their reported levels of pathogenicity.

We then selected four publicly available binary-classification models that are capable of training on imbalanced datasets: the `RandomForest` model by *sklearn* (Pedregosa *et al.*, 2011), the `LogisticRegression` model by *sklearn*, the `BalancedRandomForest` model by *imblearn* (Lematre *et al.*, 2017), and the `EasyEnsembleClassifier` model by *imblearn*. These classifiers were chosen for three main reasons: 1) their ability to handle imbalanced data (i.e. far more unreported variants than reported variants), 2) their ability to scale to the size of the training and testing datasets, and 3) they are freely available implementations that can be tuned, trained, and tested with relative ease in the same Python framework. The two random forest classifiers build collections of decision trees that weight each training input by its class frequency. Logistic regression calculates the probability of a value belonging to a particular class, again weighting by the class frequency. In contrast to the other three tested methods, the ensemble classification balances the training input using random under-sampling and then trains an ensemble of AdaBoost learners. For more details on each classifier, please refer to the *sklearn* and *imblearn* documentations (Pedregosa *et al.*, 2011; Lematre *et al.*, 2017).

Initially, we also tested the support vector classifier by *sklearn* (`SVC`), the multi-layer perceptron by *sklearn* (`MLPClassifier`), and the random under-sampling AdaBoost classifier by *imblearn* (`RUSBoostClassifier`). Each of these was excluded from our results due to, respectively, scaling issues with the training size, failure to handle the data imbalance, and overfitting to the training set. While we did not achieve positive results using these three implementations, it may be possible to use the methods through another implementation.

For each of our tested classifiers, we selected a list of hyperparameters to test and tested each possible combination of those hyperparameters. For each classifier and set of hyperparameters, we performed stratified 10-fold cross validation on the training variants and recorded the balanced accuracy (i.e. weighted accuracy based on inverse class frequency) and the F1 scores (i.e. harmonic mean between precision and recall). For each classifier type, we saved the hyperparameters and classifier with the best average F1 score (this is recommended for imbalanced datasets). These four tuned classifiers were then trained on the full training set and tested against the unseen set of test proband cases. The set of hyperparameters tested along with the highest performance setting for each hyperparameter can be found in the Supplementary Documents.

## 4 Results

### 4.1 Classifier Statistics

The hyperparameters for each classifier were tuned using 10-fold cross validation and the resulting average and standard deviation of balanced accuracy is reported in Table 2. After fitting the tuned classifiers to the full training set, we evaluated the classifiers on the testing set by calculating the area under the receiver operator curve (AUROC) and area under the precision-recall curve (AUPRC) (also shown in Table 2). Figure 1 shows the corresponding receiver operator curves and precision-recall curves for the results from the testing set on all four classifiers.

**Table 2:** Classifier performance statistics. For each tuned classifier, we show performance measures commonly used for classifiers (from left to right): 10-fold cross validation balanced accuracy (CV10 Acc.), area under the receiver operator curve (AUROC), and area under the precision-recall curve (AUPRC). The CV10 Acc. was gathered during hyperparameter tuning by calculating the average and standard deviation of the 10-fold cross validation. AUROC and AUPRC was evaluated on the testing set after hyperparameter tuning and fitting to the full training set.

| Classifier | CV10 Acc. | AUROC | AUPRC |
| --- | --- | --- | --- |
| RandomForest(sklearn) | 0.84+-0.13 | 0.9282 | 0.1961 |
| LogisticRegression(sklearn) | 0.84+-0.13 | 0.9300 | 0.2458 |
| BalancedRandomForest(imblearn) | 0.86+-0.11 | 0.9313 | 0.2015 |
| EasyEnsembleClassifier(imblearn) | 0.85+-0.08 | 0.9303 | 0.1918 |

**Figure 1:**
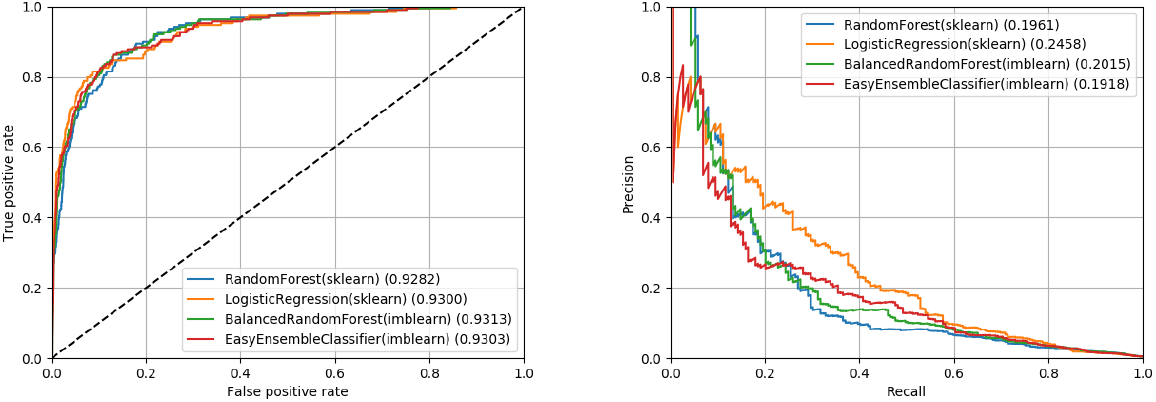
Receiver operator and precision-recall curves. These figures show the performance of the four classifiers on the testing set after hyperparameter tuning and fitting to the training set. On the left, we show the receiver operator curve (false positive rate against the true positive rate). On the right, we show the precision recall curve. Area under the curve (AUROC or AUPRC) is reported beside each method in the legend.

From these metrics, we can see that all four classifiers have a similar performance with regards to AUROC. However, all classifiers have a relatively poor performance from a precision-recall perspective (best AUPRC was only 0.2458). This indicates that from a classification perspective, these classifiers would identify a high number of false positives relative to the true positives unless a very conservative cutoff score was used.

### 4.2 Ranking Statistics

In addition to the classifier performance statistics, we also quantified the performance of each classifier as a ranking system. For each proband, we calculated the probability of each class (reported or not reported) for each variant and ordered them from highest to lowest probability of being reported. We then calculated median and mean rank statistics for the reported variants. Additionally, we quantified the percentage of reported variants that were ranked in the top 1, 10, and 20 variants in each case. While the classifiers were trained as a binary classification system, we broke down the results further to demonstrate differences between variants that were clinically reported as a variant of uncertain significance (VUS), likely pathogenic, and pathogenic.

For comparison, we selected to run Exomiser (Smedley *et al.*, 2015a), Phen-Gen (Javed *et al.*, 2014), and DeepPVP (Boudellioua *et al.*, 2019). For each tool, we input the exact same set of phenotype terms used by the classifiers we tested. Additionally, we used the same set of pre-filtered variants from Codicem as input to each ranking algorithm. As a result, all external tools and our trained classifiers are ranking on identical phenotype and variant information.

For Exomiser, we followed the installation on their website to install Exomiser CLI v.11.0.0 along with version 1811 for hg19 data sources. We ran Exomiser twice, once using the default hiPhive prioritizer (incorporates knowledge from human, mouse, and fish) and once using the human only version of the hiPhive prioritizer (this was recommended instead of the PhenIX algorithm (Zemojtel *et al.*, 2014)). Phen-Gen V1 was run using the pre-compiled binary using the “dominant” and “genomic” modes to maximize the output. Of note, Phen-Gen was the only external method that did not fully rank all variants, so we conservatively assumed that any absent variants were at the next best possible rank. Thus, the reported Phen-Gen comparisons are an optimistic representation for this test data. Finally, DeepPVP v2.1 was run using the instructions available on their website. Details on the exact installation and execution for each external tool can be found in the Supplementary Documents.

Finally, we added two control scores for comparison: CADD scaled and HPO-cosine. These scores were inputs to each classifier, but also represent two common ways one might naively order variants after filtering (by predicted deleteriousness and by similarity to phenotype). The results for the two control scores, all four external tools, and all four trained classifiers are shown in Table 3.

**Table 3:** Ranking performance statistics. This table shows the ranking performance statistics for all methods evaluated on our test set. CADD Scaled and HPO-cosine are single value measures that were used as inputs to the classifiers we tested. The middle four rows (two Exomiser runs, Phen-Gen, and DeepPVP) represent external tools that ranked the same set of variants as the classifier algorithms. Phen-Gen was the only external tool that did not rank every variant in the set, so we conservatively assumed unranked variants were at the next best position despite being unranked. The bottom four rows are the tuned, binary classification methods tested in this paper. The “Case Rank” columns show the median and mean ranks for all reported variants along with the variants split into their reported pathogenicity (variant of uncertain significance (VUS), likely pathogenic (LP), or pathogenic (Path.)) derived from the ACMG guidelines. The “Percentage in Top X Variants” columns show the percentage of variants that were found in the top 1, 10, and 20 variants in a case after ranking by the corresponding method. All values in this table were generated using only the Codicem-filtered variants from testing set.

| Ranking System | Case Rank - Median (Mean) |  |  |  | Percentage in Top X Variants - X=(1, 10, 20) |  |  |  |
| --- | --- | --- | --- | --- | --- | --- | --- | --- |
|  | All (n=189) | VUS (n=111) | LP (n=42) | Path. (n=36) | All (n=189) | VUS (n=111) | LP (n=42) | Path. (n=36) |
| CADD Scaled | 57.0 (99.13) | 69.0 (107.78) | 39.5 (91.24) | 28.0 (81.67) | 4, 17, 24 | 0, 9, 15 | 7, 21, 30 | 13, 41, 47 |
| HPO-cosine | 22.0 (53.96) | 22.0 (56.05) | 26.0 (56.38) | 19.5 (44.69) | 7, 32, 47 | 7, 31, 48 | 7, 28, 40 | 8, 38, 50 |
| Exomiser(hiPhive) | 79.0 (105.34) | 85.0 (116.33) | 93.5 (101.10) | 34.0 (76.42) | 7, 29, 36 | 6, 30, 36 | 2, 16, 28 | 16, 38, 44 |
| Exomiser(hiPhive, human only) | 35.0 (53.60) | 37.0 (63.84) | 34.0 (45.60) | 24.5 (31.36) | 7, 28, 37 | 6, 28, 36 | 2, 16, 30 | 16, 38, 50 |
| Phen-Gen | 55.0 (48.66) | 65.0 (52.91) | 47.0 (47.48) | 24.0 (36.92) | 4, 21, 30 | 5, 20, 27 | 4, 16, 26 | 2, 27, 44 |
| DeepPVP | 15.0 (76.95) | 23.0 (79.68) | 19.5 (84.95) | 6.0 (59.19) | 11, 42, 52 | 4, 36, 47 | 16, 42, 50 | 27, 61, 72 |
| RandomForest(sklearn) | 10.0 (29.64) | 15.0 (39.27) | 8.0 (20.07) | 4.0 (11.11) | 16, 53, 65 | 9, 45, 55 | 19, 61, 76 | 36, 69, 80 |
| LogisticRegression(sklearn) | 6.0 (29.23) | 14.0 (39.86) | 3.0 (22.05) | 1.0 (4.83) | 23, 58, 72 | 13, 44, 62 | 26, 71, 80 | 52, 88, 94 |
| BalancedRandomForest(imblearn) | 8.0 (28.24) | 14.0 (38.64) | 5.0 (17.67) | 3.0 (8.50) | 16, 55, 67 | 9, 44, 57 | 23, 66, 76 | 33, 77, 86 |
| EasyEnsembleClassifier(imblearn) | 7.0 (28.72) | 15.0 (40.15) | 6.0 (18.40) | 2.0 (5.50) | 17, 58, 70 | 12, 43, 60 | 14, 71, 78 | 36, 88, 94 |

In the overall data, all four classifiers outperform the single-value measures and external tools across the board. As one would intuitively expect, all classifiers perform better as the returned pathogenicity increases ranking 33-52% of pathogenic variants in the first position and 80-94% of pathogenic variants in the top 20.

## 5 Conclusion

We assessed the application of binary classification algorithms for identifying variants that were ultimately reported on a clinical report for rare disease patients. We trained and tested these algorithms using real patient variants and phenotype terms obtained from the Undiagnosed Diseases Network (UDN). From a classification perspective, we found that these methods tend to have low precision scores, meaning a high number of false positives were identified by each method. However, when evaluated as a ranking system, all four methods out-performed the single-measure ranking systems and external tools that were tested. The classifiers had median ranks of 6.0-10.0 for all reported variants and ranked 65-72% of those variants in the top 20 for the case. For “Pathogenic” variants, the median ranks were 1.0-4.0 and 80-94% of those variants were ranked in the top 20 for the case. While these algorithms are not perfect classifiers, their use as a prioritization system is quite promising.

There are two major factors that we believe is influencing the classifiers’ performance relative to the externally tested tools. First, all results were generated using real-world patients from the UDN, but only our four classifiers were trained on real-world patients from the UDN. In contrast, the four external tools were primarily evaluated and/or trained using simulations that do not capture the variation and/or noise that is apparent in UDN patients. Second, the four classifiers we tested have far more information (i.e. features) available to them than the external tools. As noted in our methods, we tried to reflect an analyst’s view of each variant as much as possible, starting with 95 features that were pruned down to 20 features used by each classifier. Incorporating the same set of features and/or training on real-world patients may improve the externally tested tools with respect to these classifiers.

We expect these classification algorithms could be refined in a variety of ways. First, adding new features could lead to increased performance in the classifiers. Additionally, some of the features represent data that is not freely available to the research community, so replacing those features with publicly accessible sources would likely influence the results. Second, there may be a better classification algorithms for this type of data. The four selected classifiers were all freely available methods intended to handle the large class imbalance in the training set, but other algorithms that aren’t as readily available may have better performance. Finally, training the classifier on different patient populations will likely yield different results, especially in terms of feature selection and feature importances.

Overall, we believe the classifiers trained in VarSight represent a significant step forward in tackling real clinical data. The tested classifiers improved our ability to prioritize variants despite the variability and uncertainty injected by real-world patients. Ultimately, we believe implementing these classifiers will enable analysts to assess the best candidate variants first, allowing for faster clinical throughput and increased automation in the future.

## Acknowledgements

We are grateful for the participation of patients and family members of the UDN and all of their referring clinicians. We would like to acknowledge all of the teams within the UDN including the coordinating center and all of the clinical sites working hard to provide definitive diagnoses for UDN patients. We would also like to acknowledge the UDN whole genome sequencing core headed by Dr. Shawn Levy.

## Funding

This work was supported in part by the Intramural Research Program of the National Human Genome Research Institute and the NIH Common Fund through the Office of Strategic Coordination and Office of the NIH Director. Research reported in this manuscript was supported by the NIH Common Fund through the Office of Strategic Coordination and Office of the NIH Director under award numbers U01HG007530, U01HG007674, U01HG007703, U01HG007709, U01HG007672, U01HG007690, U01HG007708, U01HG007942, U01HG007943, U54NS093793, and U01TR001395. The content is solely the responsibility of the authors and does not necessarily represent the official views of the NIH.

## References

Adzhubei, Ivan, Daniel M. Jordan, and Shamil R. Sunyaev. “Predicting functional effect of human missense mutations using PolyPhen?2.” Current protocols in human genetics 76.1 (2013): 7–20.

Bagnall, Richard D., et al. “Whole genome sequencing improves outcomes of genetic testing in patients with hypertrophic cardiomyopathy.” Journal of the American College of Cardiology 72.4 (2018): 419–429.

Bick, David, et al. “Successful application of whole genome sequencing in a medical genetics clinic.” Journal of pediatric genetics 6.02 (2017): 061–076.

Boudellioua, Imane, et al. “DeepPVP: phenotype-based prioritization of causative variants using deep learning.” BMC bioinformatics 20.1 (2019): 65.

Choi, Yongwook. “A fast computation of pairwise sequence alignment scores between a protein and a set of single-locus variants of another protein.” Proceedings of the ACM Conference on Bioinformatics, Computational Biology and Biomedicine. ACM, 2012.

Cooper, Gregory M., et al. “Distribution and intensity of constraint in mammalian genomic sequence.” Genome research 15.7 (2005): 901–913.

Cornish, Adam, and Chittibabu Guda. “A comparison of variant calling pipelines using genome in a bottle as a reference.” BioMed research international 2015 (2015).

DePristo, Mark A., et al. “A framework for variation discovery and genotyping using next-generation DNA sequencing data.” Nature genetics 43.5 (2011): 491.

Desvignes, Jean-Pierre, et al. “VarAFT: a variant annotation and filtration system for human next generation sequencing data.” Nucleic acids research (2018).

Dong, Chengliang, et al. “Comparison and integration of deleteriousness prediction methods for nonsynonymous SNVs in whole exome sequencing studies.” Human molecular genetics 24.8 (2014): 2125–2137.

Envision Genomics. “Codicem Analysis Platform.” Envision Genomics. URL: http://envisiongenomics.com/codicem-analysis-platform/.

Girdea, Marta, et al. “PhenoTips: Patient Phenotyping Software for Clinical and Research Use.” Human mutation 34.8 (2013): 1057–1065.

Hamosh, Ada, et al. “Online Mendelian Inheritance in Man (OMIM), a knowledgebase of human genes and genetic disorders.” Nucleic acids research 33.suppl_1 (2005): D514–D517.

Hu, Hao, et al. “VAAST 2.0: improved variant classification and disease-gene identification using a conservation-controlled amino acid substitution matrix.” Genetic epidemiology 37.6 (2013): 622–634.

Huang, Ni, et al. “Characterising and predicting haploinsufficiency in the human genome.” PLoS genetics 6.10 (2010): e1001154.

Jger, Marten, et al. “Jannovar: A Java Library for Exome Annotation.” Human mutation 35.5 (2014): 548–555.

Javed, Asif, Saloni Agrawal, and Pauline C. Ng. “Phen-Gen: combining phenotype and genotype to analyze rare disorders.” Nature methods 11.9 (2014): 935.

Jian, Xueqiu, Eric Boerwinkle, and Xiaoming Liu. “In silico prediction of splice-altering single nucleotide variants in the human genome.” Nucleic acids research 42.22 (2014): 13534–13544.

Khler, Sebastian, et al. “Clinical diagnostics in human genetics with semantic similarity searches in ontologies.” The American Journal of Human Genetics 85.4 (2009): 457–464.

Koehler, Sebastian. “Ontology-based similarity calculations with an improved annotation model.” bioRxiv (2017): 199554.

Khler, Sebastian, et al. “Expansion of the Human Phenotype Ontology (HPO) knowledge base and resources.” Nucleic acids research (2018).

Kumar, Prateek, Steven Henikoff, and Pauline C. Ng. “Predicting the effects of coding non-synonymous variants on protein function using the SIFT algorithm.” Nature protocols 4.7 (2009): 1073.

Landrum, Melissa J., et al. “ClinVar: public archive of interpretations of clinically relevant variants.” Nucleic acids research 44.D1 (2015): D862–D868.

Lek, Monkol, et al. “Analysis of protein-coding genetic variation in 60,706 humans.” Nature 536.7616 (2016): 285.

Lematre, Guillaume, Fernando Nogueira, and Christos K. Aridas. “Imbalanced-learn: A python toolbox to tackle the curse of imbalanced datasets in machine learning.” The Journal of Machine Learning Research 18.1 (2017): 559–563.

Li, Heng. “Aligning sequence reads, clone sequences and assembly contigs with BWA-MEM.” arXiv preprint arXiv:1303.3997 (2013).

Page, Lawrence, et al. “The PageRank citation ranking: Bringing order to the web.” Stanford InfoLab, 1999.

Pedregosa, Fabian, et al. “Scikit-learn: Machine learning in Python.” Journal of machine learning research 12.Oct (2011): 2825–2830.

Petrovski, Slav, et al. “Genic intolerance to functional variation and the interpretation of personal genomes.” PLoS genetics 9.8 (2013): e1003709.

Ramoni, Rachel B. et al. “The undiagnosed diseases network: accelerating discovery about health and disease.” The American Journal of Human Genetics 100.2 (2017): 185–192.

Rao, Aditya, et al. “Phenotype-driven gene prioritization for rare diseases using graph convolution on heterogeneous networks.” BMC medical genomics 11.1 (2018): 57.

Rehm, Heidi L., et al. “ACMG clinical laboratory standards for next-generation sequencing.” Genetics in medicine 15.9 (2013): 733.

Rentzsch, Philipp, et al. “CADD: predicting the deleteriousness of variants throughout the human genome.” Nucleic acids research (2018).

Richards, Sue, et al. “Standards and guidelines for the interpretation of sequence variants: a joint consensus recommendation of the American College of Medical Genetics and Genomics and the Association for Molecular Pathology.” Genetics in medicine 17.5 (2015): 405.

Roy, Somak, et al. “Standards and guidelines for validating next-generation sequencing bioinformatics pipelines: a joint recommendation of the Association for Molecular Pathology and the College of American Pathologists.” The Journal of Molecular Diagnostics 20.1 (2018): 4–27.

Siepel, Adam, Katherine S. Pollard, and David Haussler. “New methods for detecting lineage-specific selection.” Annual International Conference on Research in Computational Molecular Biology. Springer, Berlin, Heidelberg, 2006.

Singleton, Marc V., et al. “Phevor combines multiple biomedical ontologies for accurate identification of disease-causing alleles in single individuals and small nuclear families.” The American Journal of Human Genetics 94.4 (2014): 599–610.

Smedley, Damian, et al. “Next-generation diagnostics and disease-gene discovery with the Exomiser.” Nature protocols 10.12 (2015): 2004.

Smedley, Damian, and Peter N. Robinson. “Phenotype-driven strategies for exome prioritization of human Mendelian disease genes.” Genome medicine 7.1 (2015): 81.

Steinberg, Julia, et al. “Haploinsufficiency predictions without study bias.” Nucleic acids research 43.15 (2015): e101–e101.

Stenson, Peter D., et al. “Human gene mutation database (HGMD): 2003 update.” Human mutation 21.6 (2003): 577–581.

Sweeney, Nathaly M., et al. “The case for early use of rapid whole genome sequencing in management of critically ill infants: Late diagnosis of Coffin-Siris syndrome in an infant with left congenital diaphragmatic hernia, congenital heart disease and recurrent infections.” Molecular Case Studies (2018): mcs-a002469.

Wang, Kai, Mingyao Li, and Hakon Hakonarson. “ANNOVAR: functional annotation of genetic variants from high-throughput sequencing data.” Nucleic acids research 38.16 (2010): e164–e164.

Wei, Chih-Hsuan, Hung-Yu Kao, and Zhiyong Lu. “PubTator: a web-based text mining tool for assisting biocuration.” Nucleic acids research 41.W1 (2013): W518–W522.

Wilk, Brandon, James M. Holt, and Elizabeth A. Worthey. “PyxisMap.” HudsonAlpha Institute for Biotechnology. URL: https://github.com/HudsonAlpha/LayeredGraph.

Worthey, Elizabeth A. “Analysis and Annotation of Whole-Genome or Whole-Exome Sequencing Derived Variants for Clinical Diagnosis.” Current protocols in human genetics 95.1 (2017): 9–24.

Yang, Hui, Peter N. Robinson, and Kai Wang. “Phenolyzer: phenotype-based prioritization of candidate genes for human diseases.” Nature methods 12.9 (2015): 841.

Zemojtel, Tomasz, et al. “Effective diagnosis of genetic disease by computational phenotype analysis of the disease-associated genome.” Science translational medicine 6.252 (2014): 252ra123–252ra123.

