## Supplemental document for "VarSight: Prioritizing Clinically Reported Variants with Binary Classification Algorithms"

### Supplementary Document

James M. Holt *et al.*

March 27, 2019

### 1 Model Features

#### 1.1 Overview

There are a total of 50 features used as inputs to the models. Two of those features are phenotype-based features and the remaining 48 are derived from Codicem (`http://envisiongenomics.com/codicem-analysis-platform/`) extractions.

#### 1.2 Phenotype Features

For both methods, we relied on annotations and ontological structures provided by the Human Phenotype Ontology (HPO). Specifically, we used the `hp.obo` and `ALL_SOURCES_ALL_FREQUENCIES_phenotype_to_genes.txt` corresponding to data version `releases/2018-10-09`.

##### 1.2.1 HPO-cosine

The first phenotype feature is called the "HPO-cosine" score. Given $P$ terms in the HPO, a phenotype profile is represented by an $P$-dimensional vector where dimension $p$ is set to 1.0 if the corresponding phenotype term is present in the profile. Note that if a phenotype is present in the profile, all ancestor terms as defined by the ontology are automatically set as present as well. Given two profiles, we calculate the "HPO-cosine" score by performing the dot product calculation on the vectors and normalizing by their lengths. Using this measure, values close to 1.0 are highly similar (the vectors are "pointing" in similar directions) whereas those close to 0.0 are not (the vectors are mostly perpendicular).

For each gene, we created a phenotype profile based on the HPO annotations relating genes to phenotypes. For each patient, we created a phenotype profile and calculated the cosine score between the patient profile and all genes to create an ordered rank from 1 to $G$, where $G$ is the total number of genes annotated by the HPO. For each variant, we then found the smallest rank, $R$, of genes tied to the variant and stored the normalized value ($\frac{R}{G}$) as the "HPO-cosine" feature tied to the variant. For precise implementation details, please refer to the source code (`https://github.com/HudsonAlpha/VarSight/blob/master/scripts/HPOUtil.py`).

##### 1.2.2 PyxisMap

The second phenotype is derived from PyxisMap (`https://github.com/HudsonAlpha/LayeredGraph`). We derived our data from PyxisMap v1.2 and we ran the standard installation script that downloads all required data sources on December 19, 2018.

To run PyxisMap, we passed in the patient phenotype terms and obtained an ordered rank of all genes. For each variant, we then selected the best rank from genes tied to the variant, normalized the rank, and stored it as the "PyxisMap" feature tied to the variant. Note that this process is identical to how ranks were handled in the "HPO-cosine" feature. For precise implementation details, please refer to the source code (`https://github.com/HudsonAlpha/VarSight/blob/master/scripts/PyxisMapUtil.py`).

#### 1.3 Codicem Features

Codicem annotates each variant using many different data sources. We selected the annotations that we believed were most frequently used by analysts when reviewing a case. Each annotation is assigned a feature type corresponding to how that annotation was turned into a feature for the models:

1. float - The feature is a single value floating-point decimal that is copied from the annotation.

2. single - The annotation is a categorical field with a single value per variant. Each allowed value is assigned a numerical value, and that value is used as the feature.

3. multiple - The annotation is a categorical field with multiple values per variant. The total number of occurrences of each allowed value is calculated per variant and each is stored as a feature. This is the only annotation type resulting in multiple features per variant.

4. float_reduce - The annotation is a floating-point decimal with multiple values per variant. These float values are combined using a reduction method (typically, minimum or maximum as defined by the JSON configuration) and the single-value result of this reduction is used as the feature.

The JSON describing features is available on GitHub (`https://github.com/HudsonAlpha/VarSight/blob/master/CODI_metadata/fields_metadata.json`), and a selection of that information is reproduced here. Table 1 contains a list of each Codicem annotations that was automatically generated from the JSON.

| Feature Label | Feature Type | Description |
|---|---|---|
| percent reads | float | The percentage of reads containing the alternate allele at a variant site. |
| total depth | float | The total number of reads covering a variant site. |
| CADD Scaled | float | The phred-scaled version of the CADD score that estimates the pathogenicity of a variant in the human genome. |
| phylop conservation | float | The PhyloP conservation score is a measure of evolutionary conservation at an individual genomic site. This value is the 46-way PhyloP conservation score that uses the alignments of 45 vertebrate species to the human genome. |

| **Feature Label** | **Feature Type** | **Description** |
|---|---|---|
| phylop100 conservation | float | The PhyloP conservation score is a measure of evolutionary conservation at an individual genomic site. This value is the 100-way PhyloP conservation score that uses the alignments of 99 vertebrate species to the human genome. |
| phastcon100 conservation | float | The PhastCon conservation score is a measure of evolutionary conservation that factors in surrounding nucleotides (as opposed to a single base). This value is the 100-way PhastCon conservation score that uses the alignments of 99 vertebrate species to the human genome. |
| Mappability | float | Mappability is a measure of how unique a location in the genome is based on the surrounding 100bp region. Low mappability may cause issues during alignment and lead to false positives/negatives during variant calling. |
| GERP rsScore | float | The GERP rejected substitutions score identifies sites in the genome that are under some functional constraint measured by the absence of substitutions at the site. |
| Gnomad Exome AF | float | GnomAD is a population frequency database. This value corresponds to the allele frequency in the exome portion of the dataset. |
| Gnomad Exome Hom alt allele count | float | GnomAD is a population frequency database. This value corresponds to the number of homozygous alternate calls in the exome portion of the dataset. |
| Gnomad Exome Hemi alt allele count | float | GnomAD is a population frequency database. This value corresponds to the number of hemizygous alternate calls in the exome portion of the dataset. |
| Gnomad Exome total allele count | float | GnomAD is a population frequency database. This value corresponds to the number of alleles called in the exome portion of the dataset. |
| Gnomad Genome AF | float | GnomAD is a population frequency database. This value corresponds to the allele frequency in the genome portion of the dataset. |

| Feature Label | Feature Type | Description |
|---|---|---|
| Gnomad Genome Hom alt allele count | float | GnomAD is a population frequency database. This value corresponds to the number of homozygous alternate calls in the genome portion of the dataset. |
| Gnomad Genome Hemi alt allele count | float | GnomAD is a population frequency database. This value corresponds to the number of hemizygous alternate calls in the genome portion of the dataset. |
| Gnomad Genome total allele count | float | GnomAD is a population frequency database. This value corresponds to the number of alleles called in the genome portion of the dataset. |
| Type | single | This value stores the type of variant (SNV, insertion, or deletion). |
| HGMD assessment type | multiple | HGMD is a database storing germline mutations related to human inherited disease. This value corresponds to the assessment of how damaging the mutation was. |
| HGMD association confidence | multiple | HGMD is a database storing germline mutations related to human inherited disease. This value corresponds to how confident the assessment of the mutation was. |
| ClinVar Classification | multiple | ClinVar is a database storing relationships between human variation and phenotype. This value corresponds to the classification of how damaging a particular variant was. |
| Ensembl Regulatory Feature | multiple | Ensembl regulatory features contains information regarding if a variant is impacting annotating promoters, enhancers, etc. |
| variant_attribute | multiple | This value contains whether a variant lies in a simple repeat or other low complexity region in the genome. |
| RVIS Score | float_reduce | The Residual Variation Intolerance Score is a measure of how intolerant to variation a particular gene is in the human genome. |

| Feature Label | Feature Type | Description |
|---|---|---|
| GHIS Score | float_reduce | The Genome-wide Haplo-Insufficiency Score is a measure of gene haploinsufficiency calculated in Steinberg et al., 2015 (https://doi.org/10.1093/nar/gkv474). |
| HIS Score | float_reduce | This Haplo-Insufficiency Score is a measure of gene haploinsufficiency calculated in Huang et al., 2010 (https://doi.org/10.1371/journal.pgen.1001154). |
| Essentiality | multiple | Essentiality is a measure of how essential a gene is for humans based on comparison to orthologs in mouse defined in Georgi et al., 2013 (https://doi.org/10.1371/journal.pgen.1003484). |
| ADA Boost Splice Prediction | float_reduce | ADA Boost Splice Prediction is a measure of the impact of a variant on splicing using adaptive boosting from Jian et al., 2014 (https://doi.org/10.1093/nar/gku1206). |
| Random Forest Splice Prediction | float_reduce | Random Forest Splice Prediction is a measure of the impact of a variant on splicing using random forest from Jian et al., 2014 (https://doi.org/10.1093/nar/gku1206). |
| Meta Svm Prediction | multiple | Meta SVM is a method to predict deleteriousness of a variant in a transcript using Support Vector Machines as defined in Dong et al., 2015 (https://doi.org/10.1093/hmg/ddu733). |
| PolyPhen HV Prediction | multiple | PolyPhen predicts the impact of amino acid substitutions on a protein. This is the PolyPhen2 score based on HumVar as defined in Adzhubei et al., 2014 (https://doi.org/10.1002/0471142905.hg0720s76). |
| PolyPhen HD Prediction | multiple | PolyPhen predicts the impact of amino acid substitutions on a protein. This is the PolyPhen2 score based on HumDiv as defined in Adzhubei et al., 2014 (https://doi.org/10.1002/0471142905.hg0720s76). |

| Feature Label | Feature Type | Description |
|---|---|---|
| Provean Prediction | multiple | PROVEAN (PROtein Variant Effect ANalyzer) predicts where an amino acid substition or indel impacts biological function as defined by Choi, 2012 (`https://doi.org/10.1145/2382936.2382989`). |
| SIFT Prediction | multiple | SIFT predicts whether an amino acid substitution impacts protein function as defined by Kumar et al., 2009 (`https://doi.org/10.1038/nprot.2009.86`). |
| Effects | multiple | Effects includes the predicted impact on a transcript such as non-synonymous, possible frameshift, etc. |
| Affected Regions | multiple | Affected regions includes the regions of a transcript that are impacted by the variant such as interior coding exon, 5' UTR intron, etc. |

Table 1: Codicem feature metadata. This table shows the feature name, feature type, and a brief description of each feature that was fed into the models from Codicem.

### 2 Hyperparameter Tuning

#### 2.1 Tuning method

For each classifier, a selection of hyperparameters were tested based on recommendations from `sklearn` and `imblearn`. We performed a brute-force search over the entire hyperparameter space using the `GridSearchCV` method of `sklearn`. This method tries every combination of hyperparameters and selects the method with the best performance as defined by the user. For defining performance, we performed stratified 10-fold cross validation (`StratifiedKFold` in `sklearn`) on the training data and optimized for best F1-score (see `https://scikit-learn.org/stable/modules/generated/sklearn.metrics.f1_score.html` for more info).

#### 2.2 Tuning results

Table 2 shows the classifiers, hyperparameters, tested values, and the final tuned value for our tests.

| Classifier | Hyperparameter | Tested Values | Tuned Value |
|---|---|---|---|
| RandomForest (sklearn) | class_weight | ['balanced'] | balanced |
| | max_depth | [2, 3, 4] | 4 |
| | max_features | ['sqrt', 'log2'] | sqrt |
| | min_samples_split | [2, 3] | 2 |
| | n_estimators | [100, 200, 300] | 300 |
| | random_state | [0] | 0 |
| LogisticRegression (sklearn) | C | [0.01, 0.1, 1.0, 10.0, 100.0] | 0.01 |
| | class_weight | ['balanced'] | balanced |
| | max_iter | [200] | 200 |
| | penalty | ['l2'] | l2 |
| | random_state | [0] | 0 |
| | solver | ['newton-cg', 'liblinear'] | liblinear |
| BalancedRandomForest (imblearn) | max_depth | [2, 3, 4] | 4 |
| | max_features | ['sqrt', 'log2'] | sqrt |
| | min_samples_split | [2, 3] | 2 |
| | n_estimators | [100, 200, 300] | 200 |
| | random_state | [0] | 0 |
| EasyEnsembleClassifier (imblearn) | n_estimators | [10, 20, 30, 40, 50] | 50 |
| | random_state | [0] | 0 |

Table 2: Hyperparameters tested. Each classifier is shown on the left. For each classifier, we selected hyperparameters and test values for those parameters based on recommendations provided by the developers of each method. The selected tuned value (based on 10-fold cross validation optimizing for F1 score) for each hyperparameter is shown on the right.

### 3 External Tools

#### 3.1 Exomiser

##### 3.1.1 Installation

We followed the instructions on the Exomiser website (http://exomiser.github.io/Exomiser/manual/7/installation/) to install Exomiser CLI v.11.0.0. We downloaded the latest files for hg19 (https://data.monarchinitiative.org/exomiser/latest/1811_hg19.zip). Additionally, we downloaded data stores for ReMM v0.3.1 (https://zenodo.org/record/1197579/files/ReMM.v0.3.1.tsv.gz) and CADD v1.3 (https://krishna.gs.washington.edu/download/CADD/v1.3/whole_genome_SNVs.tsv.gz and https://krishna.gs.washington.edu/download/CADD/v1.3/InDels.tsv.gz) for use by Exomiser.

##### 3.1.2 Execution

Exomiser was run twice using two different final prioritization algorithm: hiPhive and hiPhive (human only). Here is the templated command we used to run exomiser:

```
java -Xms2g -Xmx4g -jar exomiser-cli-11.0.0.jar --analysis [ymlFN]
```

The `ymlFN` parameter is a templated file with parameters for the VCF filename, the HPO terms, and the output path. The template for the hiPhive execution is available at https://github.com/HudsonAlpha/VarSight/blob/master/ExomiserTemplates/hiPhive-template.yml and the template for the hiPhive (human only) is available at https://github.com/HudsonAlpha/VarSight/blob/master/ExomiserTemplates/hiPhive_human-template.yml. These templates were populated with the corresponding information for each patient using the same variants and HPO terms as were used in the VarSight classifiers.

#### 3.2 Phen-Gen

##### 3.2.1 Installation

We followed the instructions on the Phen-Gen website (http://phen-gen.org, click on "DOWNLOAD") to download Phen-GenV1 and followed directions in the README file to install and run Phen-Gen.

##### 3.2.2 Execution

Phen-Gen takes the HPO terms and VCF file as input to the algorithm. For the HPO terms, we created a plain-text file with one HPO term per line as instructed. We used the same pre-filtered VCF file as was used in all other external tools.

Phen-Gen also requires extra parameters that were not need by other tools, specifically requiring a pedigree file with sex information, mode of inheritance, and the predictor type. For the pedigree file, we simply created a one-member pedigree with the corresponding sex field populated with the annotated sex of the patient. For mode of inheritance, we selected dominant as this tended to rank more variants than recessive. Finally, we selected the predictor type to be "genomic" as opposed to "coding" only. Here is the templated command we used to run Phen-Gen:

```
perl phen-gen.pl input_phenotype=[hpoFN] input_vcf=[vcfFN] \
    input_ped=[pedFN] inheritance=0 predictor=0
```

`hpoFN` is the filename of the HPO file, `vcfFN` is the filename of the VCF file, and `pedFN` is the filename of the pedigree file.

#### 3.3 DeepPVP

##### 3.3.1 Installation

We followed the instructions on the GitHub README (`https://github.com/bio-ontology-research-group/phenomenet-vp`) to download the distribution file for DeepPVP v2.1.1 (`https://github.com/bio-ontology-research-group/phenomenet-vp/releases/download/v2.1.1/phenomenet-vp-2.1.zip`). We downloaded all datasets required by their instructions including the v2.1 data file (`http://bio2vec.net/pvp/data-v2.1.tar.gz`), CADD v1.3 with annotations (`http://krishna.gs.washington.edu/download/CADD/v1.3/whole_genome_SNVs_inclAnno.tsv.gz`), and DANN (`https://cbcl.ics.uci.edu/public_data/DANN/data/DANN_whole_genome_SNVs.tsv.bgz`). We then ran their preprocessing of CADD script (`http://www.bio2vec.net/pvp/generate.sh`) and moved data to the correct directories.

We created a fresh installation of Python v2.7.16 and followed the instructions to `pip install -r requirements.txt` to install all required Python packages. However, we found the program was crashing due to incompatibilities between the versions of Keras and Theano. To fix the issue, we updated to Keras v2.2.4 and the issue was resolved. Due to the relatively high performance of DeepPVP compared to other external tools, we do not believe this change influenced DeepPVP's performance.

##### 3.3.2 Execution

DeepPVP requires a VCF file and the HPO terms to execute. Here is the templated command used to run DeepPVP:

```
phenomenet-vp -y [pythonPath] -p [csv-HPO] -f [vcfFN] -of [outFN]
```

`pythonPath` is the path to our fresh Python install, `csv-HPO` is a comma-separated list of HPO terms for the patient, `vcfFN` is the filename of the VCF file, and `outFN` is the filename for storing the output.

### 4 Codicem Filtering

The following sections describes the filter that was passed to Codicem for generating the test and training set variants. Figure 1 shows the sequential order of filters, and the following subsections briefly describes each filter unit's purpose.

#### 4.1 Total Depth filter

The total depth filter is primarily a QC-related filter intended to remove variant calls with total coverage. This filter only passes variants that have at least 8 reads overlapping the locus. Figure 2 shows a visualization of the filter.

#### 4.2 Percentage of Reads Supporting Allele filter.

This filter is also a QC-related filter intended to remove variant calls with low support for the alternate allele. The filter only passes variants with more than 15% of the reads supporting the alternate allele. Figure 3 shows a visualization of the filter.

#### 4.3 Population Allele Frequency filter.

This filter removes variants that have a high population frequency. There are three general ways for a variant to pass this filter: 1) be a rare variant in GnomAD AND a rare variant in ExAC; 2) be semi-rare in GnomAD, semi-rare in ExAC, AND have a high-confidence damaging annotation from HGMD; or 3) be semi-rare in GnomAD, semi-rare in ExAC, AND have a ClinVar classification labeling it as pathogenic, likely pathogenic, or a drug response. Additionally, there is one variant from the gene *F5* (rs6025) allowed through this filter. Note that this specific variant has a very high alternate allele frequency because the reference allele is actually the disease-causing mutation. Figure 4 shows a visualization of the filter.

#### 4.4 HGMD, ClinVar, CADD, and Effects filter

This filter removes variants that are not predicted or annotated to have an effect on a transcript. A variant can pass this filter in any of four ways: 1) have an HGMD accession, 2) have a ClinVar accession, 3) have an effect that is predicted to modify a transcript, or 4) have a very high CADD score. Figure 5 shows a visualization of the filter.

#### 4.5 Gene Has Associated Disease filter

This filter removes variants for which there is no annotated disease name in HGMD, OMIM, or ClinVar. If any of those annotations for the variant has a disease name, then the variant will pass this filter. Figure 6 shows a visualization of the filter.

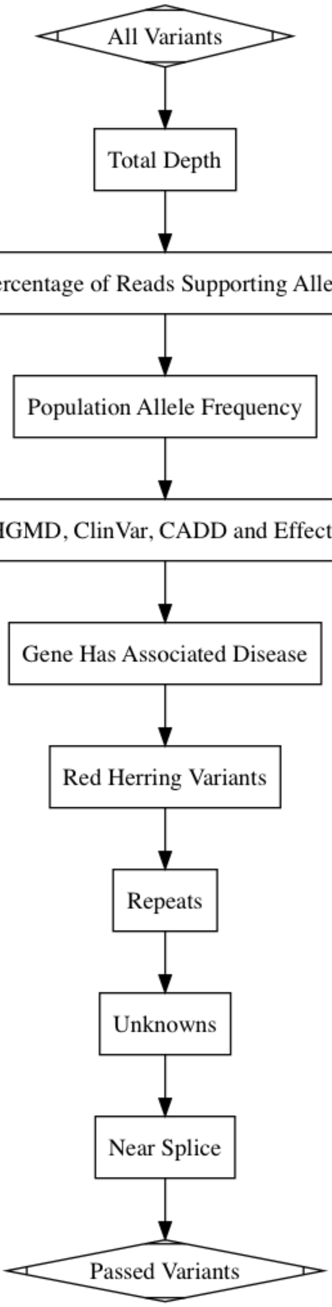

Figure 1: Filter overview. This high level image shows the sequential order of filtering, descriptions of sub-filters can be found in other sections.

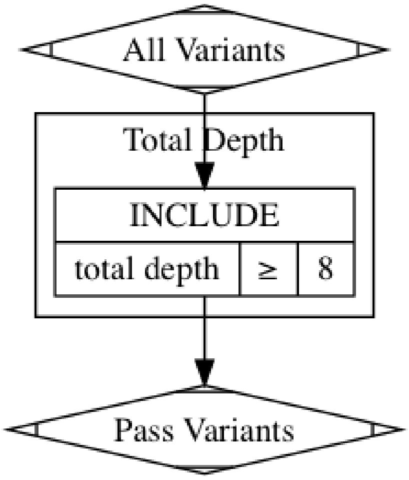

Figure 2: Total Depth filter. This filter is primarily a QC-related filter that only passes variants with at least 8 reads covering the locus.

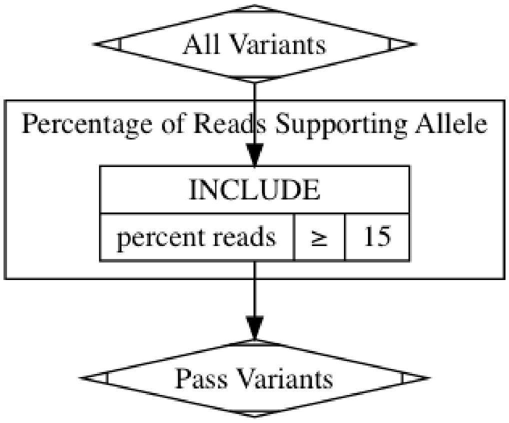

Figure 3: Percentage of Reads Supporting Allele filter. This filter is primarily a QC-related filter that only passes variants with at least 15% of the locus' reads supporting the alternate allele.

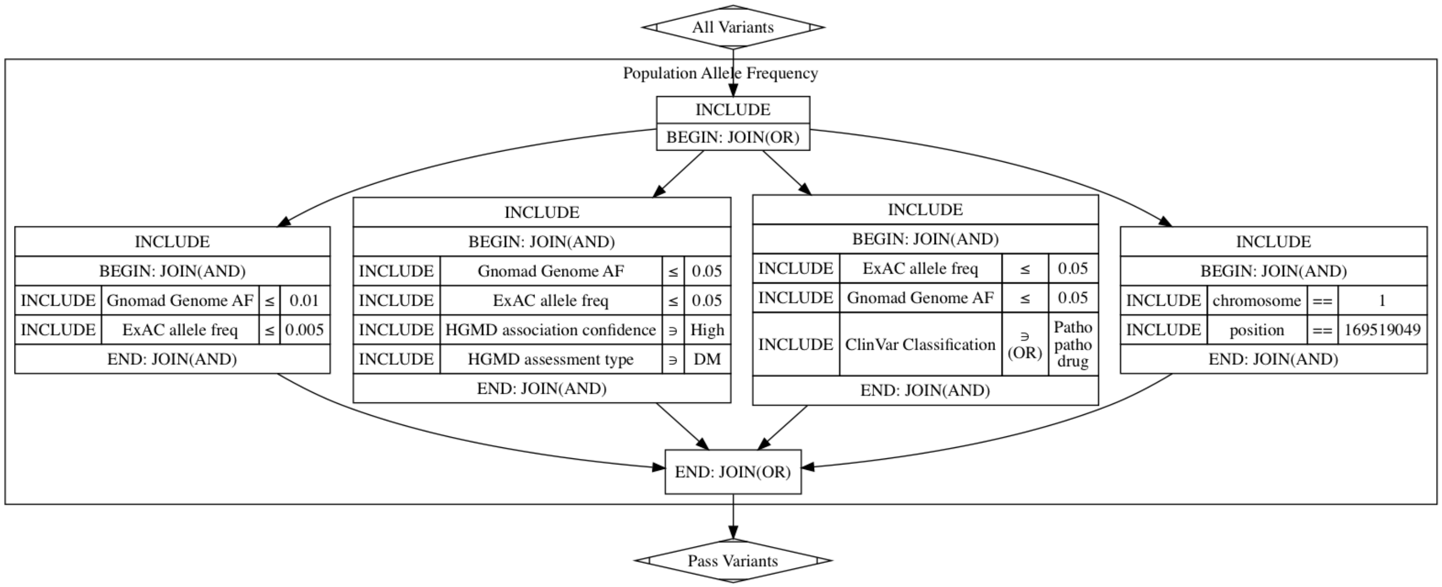

Figure 4: Population Allele Frequency filter. This filter is primarily designed to filter out variants with a high population frequency. It relies mostly on GnomAD and ExAC to determine allele frequencies. Variants that are only semi-rare are allowed through if there are appropriate HGMD or ClinVar annotations supporting them. Additionally, one variant in gene *F5* (rs6025) is allowed through because the reference allele is actually the rare, pathogenic allele.

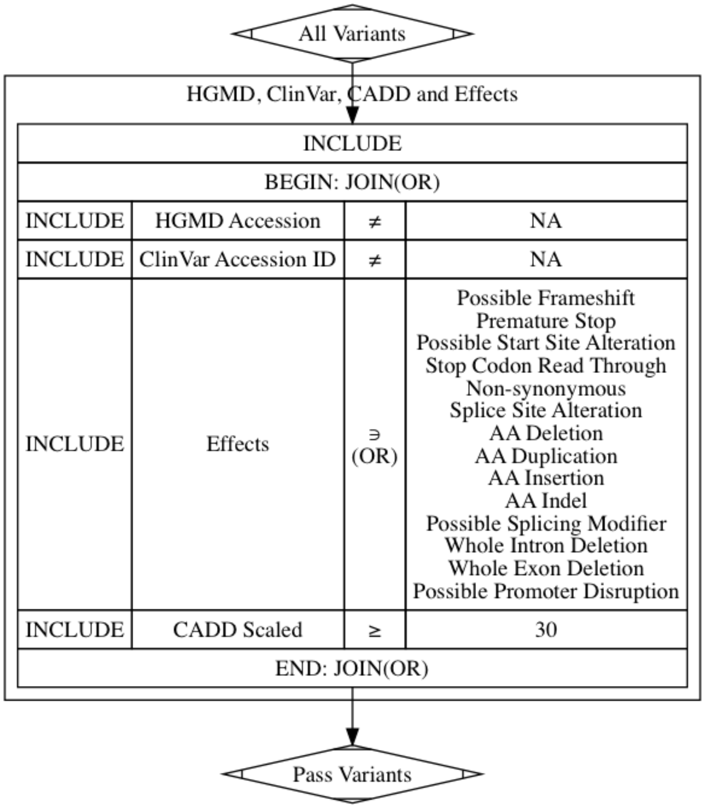

Figure 5: HGMD, ClinVar, CADD, and Effects filter. This filter is primarily designed to filter out variants that are not predicted or annotated to have an effect on a transcript.

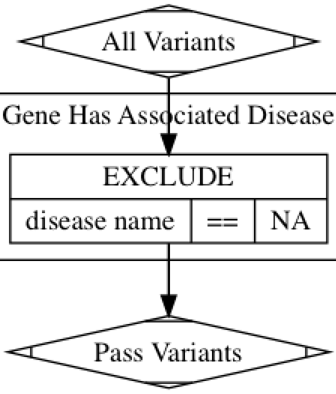

Figure 6: Gene Has Associated Disease filter. This filter removes variants for which there is no annotated disease name in HGMD, OMIM, or ClinVar.

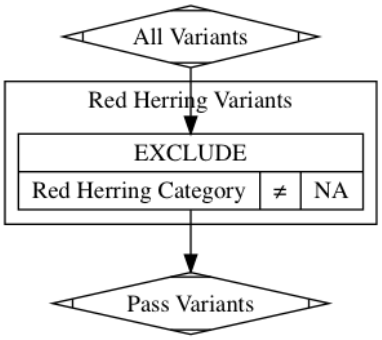

Figure 7: Red Herring filter. This filter removes variants that are commonly found through sequencing, but do not appear in population databases like gnomAD or ExAC. These variants are believed to be sequencing artifacts, so they are labeled as "Red Herring" variants and filtered out.

#### 4.6 Red Herring filter

This filter removes variants that are sequencing artifacts. Our analysts found that some variants are not found in population databases, but show up frequently (> 20% of samples) in our sequencing data. These variants are believed to be sequencing artifacts, so Codicem maintains a database of these variants to filter out during analysis. Figure 7 shows a visualization of the filter.

#### 4.7 Repeats filter

This filter removes variants that are polymorphic repeats and/or artifacts from sequencing. It filters out any variants that are found in repeat tracks if there is no associated entry in HGMD or ClinVar. Figure 8 shows a visualization of the filter.

#### 4.8 Unknowns filter

This filter removes variants that have an "Unknown" effect on transcription if there isn't additional support for that variant through either CADD or HGMD. If CADD is greater than 10 or there is an HGMD accession tied to the variant, it will pass this filter. Figure 9 shows a visualization of the filter.

#### 4.9 Near Splice filter

This filter removes near splice variants that do not have additional annotated or predicted support indicating that the variant would impact splicing. If a variant is a "Near Splice Site Alteration", that variant will only pass the filter if one of the following is true: 1) both splice predictions algorithms predict an impact

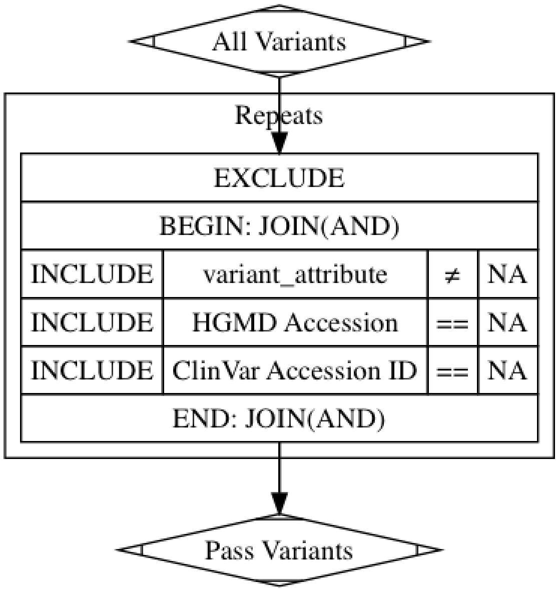

Figure 8: Repeats filter. This filter removes variants found in repeat tracks if there is no associated entry in HGMD or ClinVar. These variants are generally considered polymorphic and/or sequencing artifacts unless there are annotations from disease databases.

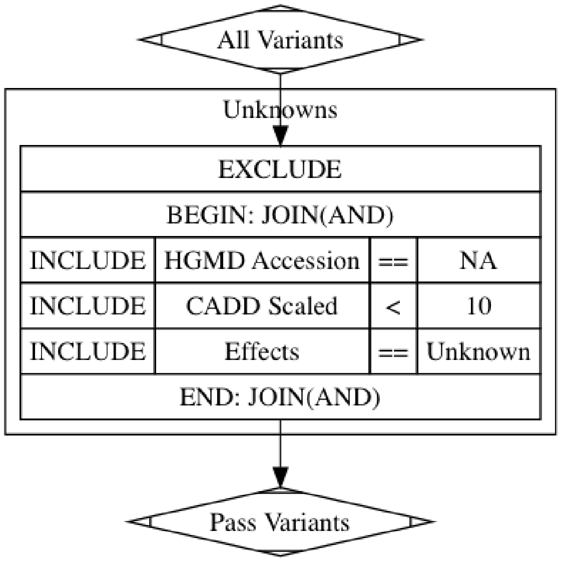

Figure 9: Unknowns filter. This filter removed variants with an “Unknown” effect on transcription if there is not additional support for that variant through either CADD or HGMD.

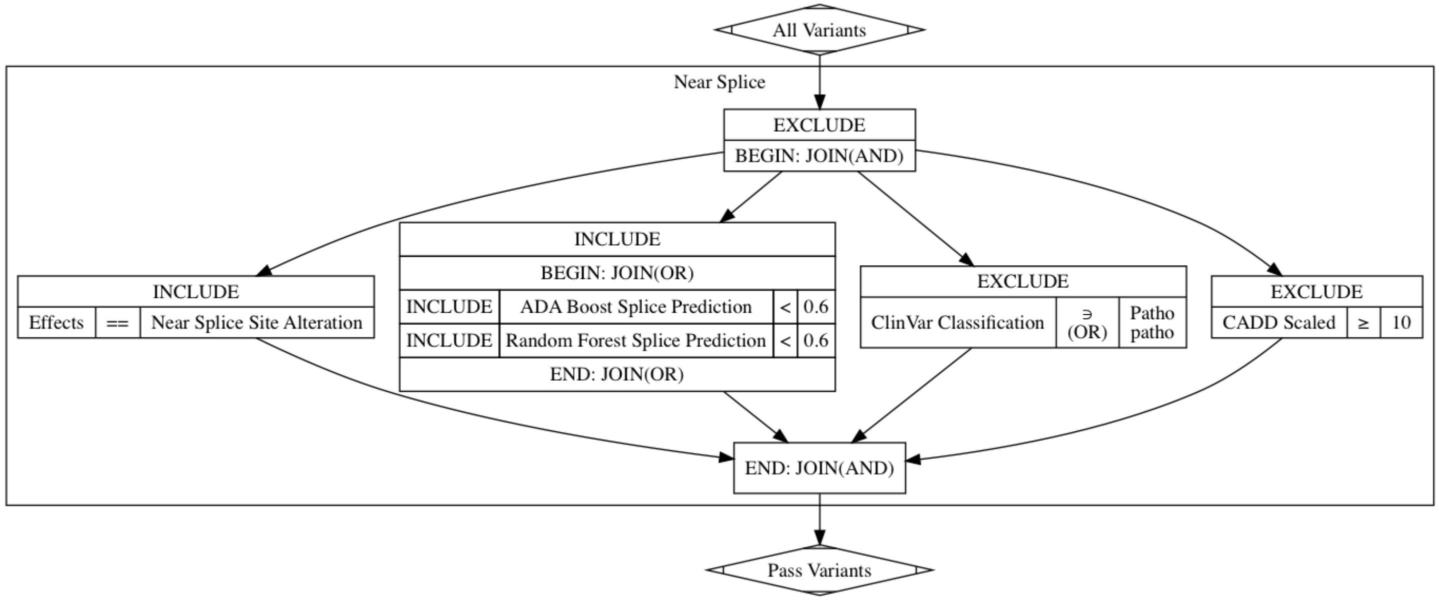

Figure 10: Near Splice filter. This filter removes “Near Splice Site Alteration” variants that are not supported to be deleterious by splice prediction algorithms, ClinVar annotations, or CADD predictions.

on splicing, 2) ClinVar has a pathogenic or likely pathogenic annotation, or 3) CADD is relatively high. Figure 10 shows a visualization of the filter.
